# LRRK2-mediated phosphorylation of HDAC6 regulates HDAC6-cytoplasmic dynein interaction and aggresome formation

**DOI:** 10.1101/554881

**Authors:** Richard M. Lucas, Claudia S. Bauer, Kavitha Chinnaiya, Aurélie Schwartzentruber, Ruby MacDonald, Mark O. Collins, Jan O. Aasly, Gunnar Brønstad, Laura Ferraiuolo, Heather Mortiboys, Kurt J. De Vos

## Abstract

Mutations in LRRK2 are the most common cause of dominantly inherited Parkinson’s disease (PD). A proportion of LRRK2 PD exhibits Lewy pathology with accumulations of α-synuclein and ubiquitin in intracellular aggregates that are indistinguishable from idiopathic PD. LRRK2 is a multi-domain protein with both GTPase and kinase activities that has been shown to affect various cellular processes including protein homeostasis, however how PD mutations in LRRK2 may lead to accumulation of ubiquitinated protein aggregates remains unclear.

A main cellular pathway to remove aggregated ubiquitinated proteins is aggrephagy: the histone deacetylase HDAC6 recognizes ubiquitinated misfolded proteins and recruits them to the molecular motor cytoplasmic dynein which transports them to the perinuclear region where they are trapped in aggresomes that are subsequently removed by macroautophagy.

Here we identified HDAC6 as a novel LRRK2 substrate and show that LRRK2 regulates HDAC6-dependent aggresome formation. LRRK2 directly interacted with the HDAC6 deacetylase domains via its Roc domain and phosphorylated HDAC6 on serine-22. Serine-22 phosphorylation of HDAC6 enhanced its interaction with cytoplasmic dynein and stimulated recruitment of ubiquitinated proteins to aggresomes. Knockdown or knockout of LRRK2 impaired HDAC6-mediated aggresome formation. PD mutant LRRK2 G2019S showed reduced interaction with HDAC6 and did not support aggresome formation to the same extend as wild type LRRK2. This was recapitulated in LRRK2 G2019S patient-derived iAstrocytes that showed an aggresome formation defect.

In conclusion our data reveal HDAC6 as a target of LRRK2 and suggest that deregulation of HDAC6-mediated aggresome formation and aggrephagy could contribute to the pathology of PD.

---

Aggresomes are pericentriolar inclusion bodies where misfolded proteins accumulate prior to removal by macroautophagy. When the ability of cells to refold proteins or to remove misfolded proteins via the ubiquitin-proteasome system (UPS) is exceeded, ubiquitinated misfolded proteins are transported by cytoplasmic dynein to aggresomes. Histone deacetylase 6 (HDAC6) plays a key role in aggresome formation. HDAC6 is a member of a family of HDACs containing 11 Zn^2+^-dependent enzymes (HDAC1–11) and 7 NAD^+^-dependent proteins (Sirtuin1–7) that are subdivided into 4 classes, Class I (HDAC1, 2, 3, and 8), Class IIa (HDAC4, 5, and 7) and IIb (HDAC6 and 10), Class III (Sirtuin1–7), and Class IV (HDAC11). HDAC6 stands out among the other HDACs because it is predominantly cytosolic, contains two catalytic domains and has a ubiquitin binding domain. Cytosolic substrates of HDAC6 include α-tubulin, Hsp90, tau, cortactin, and peroxiredoxin. In respect to aggrephagy, HDAC6 interacts with cytoplasmic dynein and recruits polyubiquitinated misfolded proteins to dynein motors for transport to aggresomes through its ubiquitin binding domain ^1^.

Lewy bodies are a hallmark pathology of Parkinson’s disease and it has been proposed that impaired handling of misfolded proteins and aggregates may contribute to their formation ^2–4^. HDAC6 is present in Lewy bodies, indicating aggrephagy may be involved in their formation ^1,5^.

LRRK2 is a multi-functional, multi-domain protein with both kinase and GTPase activity. Dominant mutations in LRRK2 are the most common cause of Parkinson’s disease, a late onset neurodegenerative movement disorder that is characterised by selective degeneration of dopaminergic neurons and the presence of intracytoplasmic proteinaceous inclusions, known as Lewy bodies (LBs) in the substantia nigra pars compacta. Pathogenic mutations have been identified in the Ras of complex proteins (Roc) GTPase protein domain (R1441C, R1441G, R1441H), the carboxy-terminal of Roc (COR) domain (Y1699C) and the kinase domain (G2019S and I2020T) ^6^. Mutations in the Roc and COR domain diminish the GTPase activity of LRRK2, whereas mutations in the kinase domain enhance its kinase activity. It is generally assumed that kinase hyperactivity is linked to neurotoxicity, but it is less clear how diminished LRRK2 GTPase activity contributes to disease ^7^.

In most cases LRRK2-associated Parkinson’s disease is clinically and pathologically indistinguishable from idiopathic late-onset PD but this may vary depending on the type of pathogenic mutations ^8^. Our previous studies have linked LRRK2 to HDAC6 and microtubule acetylation ^9^, and LRRK2 has been proposed to be involved in proteostasis and aggrephagy, but reports are conflicting and the molecular mechanisms involved poorly understood ^10–12^.

Here we investigated the role of LRRK2 in aggrephagy. We report that phosphorylation of HDAC6 by LRRK2 regulates HDAC6-dependent delivery of ubiquitinated proteins to the aggresome and show that this novel function of LRRK2 is impaired by the PD-associated G2019S mutation and in LRRK2 G2019S patient-derived iAstrocytes.

## Results

### LRRK2 kinase regulates HDAC6-dependent aggresome formation

To characterise the possible role of LRRK2 in aggresome formation, we depleted LRRK2 in HEK293 cells using siRNA and quantified aggresome formation using two well-characterised aggresome reporters, EGFP-CFTRΔF508 ^13^ and GFP-250 ^14^. The cystic fibrosis causing allele of the cystic fibrosis transmembrane conductance regulator (CFTR), ΔF508, interferes with its ability to fold; misfolded CFTRΔF508 is ubiquitinated and degraded by the proteasome. Inhibition of proteasome activity in cells expressing CFTRΔF508 causes accumulation of stable, ubiquitinated aggregates of CFTRΔF508 in aggresomes ^13^. Similar to CFTRΔF508, GFP-250, GFP fused at its COOH terminus to a 250–amino acid fragment of the Golgi protein p115, is sequestered in aggresomes upon inhibition of the proteasome. However, unlike CFTRΔF508, GFP-250 in aggresomes is not ubiquitinated ^14^.

In line with previous publications ^13,14^, in control HEK293 cells treated with non-targeting control (NTC) siRNA, both EGFP-CFTRΔF508 and GFP-250 were recruited to distinctive perinuclear inclusion bodies after treatment with MG132 to inhibit the proteasome. Co-staining with vimentin confirmed that these structures were aggresomes. SiRNA-mediated LRRK2 depletion had no effect on the formation of GFP-250 aggresomes but fully prevented EGFP-CFTRΔF508 aggresomes (Fig. 1A-B; SFig. 1). To verify the specificity of the siRNA treatment we re-expressed wild type LRRK2 and as expected this completely rescued CFTRΔF508 aggresome formation in LRRK2 siRNA treated cells (Fig. 1A). To further substantiate these findings, we turned to LRRK2 knockout (KO) MEFs ^15^, reasoning that if LRRK2 is indeed required for aggresome formation that these cells would be defective in aggresome formation. As expected, LRRK2 KO MEFS did not form EGFP-CFTRΔF508 aggresomes, and this could be rescued by expression of wild type LRRK2 (Fig. 1C). Interestingly, in LRRK2 KO MEFs we did observe perinuclear vimentin-positive spots that were reminiscent of the vimentin cages associated with aggresomes despite the absence of perinuclear EGFP-CFTRΔF508 accumulation (Fig. 1C), suggesting that loss of LRRK2 may impair recruitment of EGFP-CFTRΔF508 to aggresomes rather than the assembly of vimentin-positive aggresomes per se. This is consistent with the observation that LRRK2 was not required for formation of ubiquitin-independent GFP-250 positive aggresomes (Fig. 1B).

**Figure 1.**
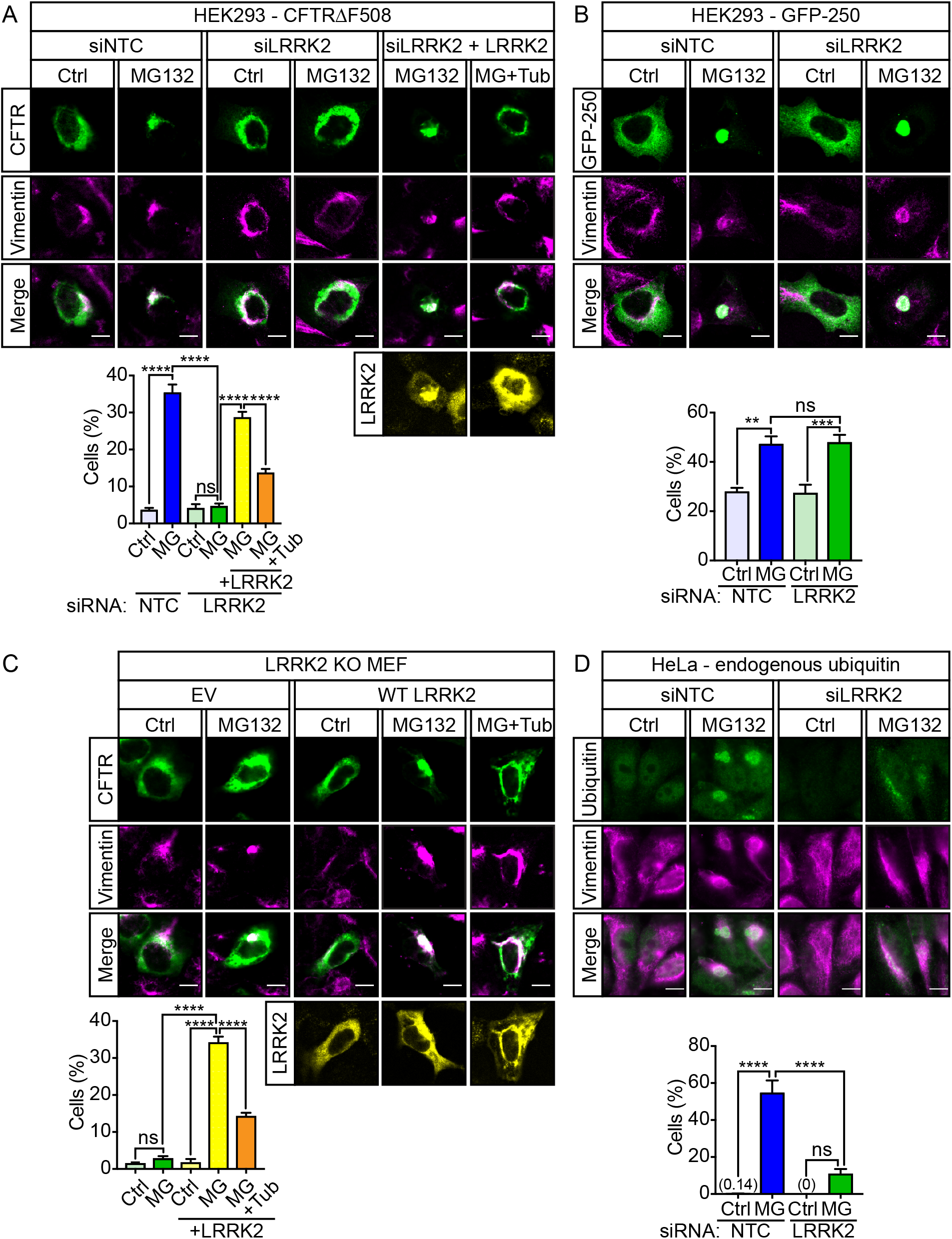
LRRK2 is required for HDAC6-dependent aggresome formation. A) Non-targeting (siNTC) or LRRK2 siRNA (siLRRK2)-treated HEK293 cells co-transfected with EGFP-CFTRΔF508 (green) and empty vector (EV) or myc-LRRK2 (yellow) were treated with vehicle, MG132 (5 µM) or MG132 + Tubastatin A (10 µM) for 4h, fixed and immunostained with an anti-vimentin antibodies (magenta). To confirm siRNA specificity cells were co-transfected with myc-LRRK2 (yellow). The percentage cells containing vimentin-positive CFTRΔF508 aggresomes was quantified (mean ± SEM; One-way ANOVA with Fisher’s LSD test, N = 9 experiments, ∼50-100 cells analysed per condition per experiment). Scale bar, 10 µm. B) Non-targeting (NTC) or LRRK2 siRNA-treated HEK293 cells transfected with GFP-250 (green) were treated with vehicle or MG132 (5 µM) for 4h and immunostained with for vimentin (magenta). The percentage cells containing vimentin-positive GFP-250 aggresomes was quantified (mean ± SEM; One-way ANOVA with Fisher’s LSD test, N = 3 experiments, ∼50-100 cells analysed per condition per experiment). Scale bar, 10 μm. C) LRRK2 KO MEFs co-transfected with EGFP-CFTRΔF508 (green) and empty vector (EV) or myc-LRRK2 (yellow) were treated with vehicle, MG132 (5 μM) or MG132 + Tubastatin A (10 µM) for 4h, fixed and immunostained with an anti-vimentin antibodies (magenta). The percentage cells containing vimentin-positive CFTRΔF508 aggresomes was quantified (mean ± SEM; One-way ANOVA with Fisher’s LSD test, N = 3 experiments, ∼50-100 cells analysed per condition per experiment). Scale bar, 10 µm. D) HeLa were treated with vehicle or MG132 (5 μM) for 18 h, fixed and immunostained for ubiquitin (green) and vimentin (magenta). The percentage cells containing ubiquitin and vimentin aggresomes was quantified (mean ± SEM; One-way ANOVA with Fisher’s LSD test, N = 3 experiments, ∼50-100 cells analysed per condition per experiment). Scale bar, 10 μm

Finally, to confirm that these observations were not just a consequence of overexpression of EGFP-CFTRΔF508 we inhibited the proteasome in NTC or LRRK2 siRNA treated HeLa cells and observed the formation of endogenous aggresomes using anti-ubiquitin antibodies. In NTC siRNA-treated HeLa accumulations of ubiquitin in the perinuclear region were readily observed after proteasome inhibition and these accumulations stained positive for vimentin, identifying them as aggresomes (Fig. 1D). In contrast, after proteasome inhibition in LRRK2 siRNA-treated HeLa, ubiquitin-positive spots were observed throughout the cytoplasm and did not accumulate in the perinuclear region, consistent with a failure to recruit ubiquitinated proteins into aggresomes (Fig. 1D).

Ubiquitinated, misfolded proteins such as CFTRΔF508 are recruited to the aggresome by HDAC6 whereas the recruitment of non-ubiquitinated proteins such as GFP-250 is HDAC6 independent ^1^. HDAC6-dependent aggresome formation requires both HDAC6 deacetylase and ubiquitin binding activity ^1^. The data above suggested a role for LRRK2 in HDAC6-dependent recruitment of ubiquitinated proteins to aggresomes. To begin to investigate this, we inhibited HDAC6 using the highly selective inhibitor Tubastatin A ^16^ in LRRK2 siRNA-treated HEK293 cells and LRRK2 KO MEFs that were reconstituted with wild type LRRK2. Tubastatin A treatment prevented the rescue of LRRK2 siRNA and KO by expression of LRRK2, showing that LRRK2 requires HDAC6 to support aggresome formation (Fig 1A, C).

### HDAC6 is a LRRK2 substrate

The data above strongly suggested that LRRK2 regulates HDAC6’s function in aggresome formation. To further investigate this, we checked if LRRK2 and HDAC6 may interact and if HDAC6 is a substrate of LRRK2.

We first investigated the possible interaction of LRRK2 and HDAC6 in co-immunoprecipitation assays in HEK293 cells. Untagged HDAC6 co-immunoprecipitated with myc-tagged LRRK2 (myc-LRRK2) from cells co-transfected with HDAC6 and myc-LRRK2, but not from HDAC6 or myc-LRRK2 only transfected cells (Fig. 2A).

**Figure 2.**
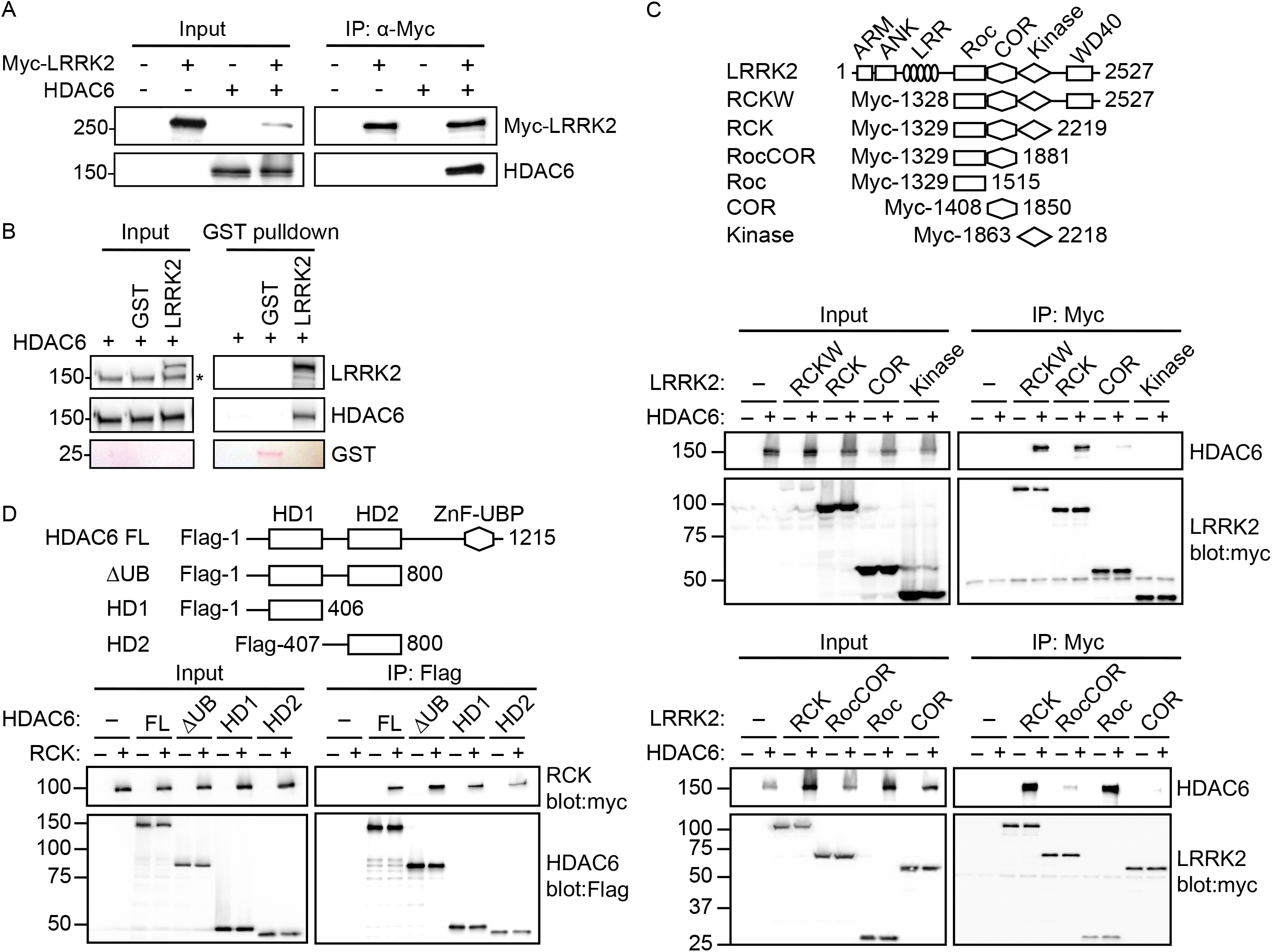
LRRK2 interacts with HDAC6. A) Myc-LRRK2 was immunoprecipitated from cell lysates made from HEK293 cells co-transfected with empty vector (–) and myc-LRRK2 or HDAC6, or myc-LRRK2 + HDAC6, as indicated. Input and immune pellets were probed with anti-myc and anti-HDAC6 antibodies. B) Full-length human His-HDAC6 (1 μg) was incubated with GST-tagged human LRRK2 (aa 970-2527; 1 μg) or GST protein (1 μg). GSH beads were used to isolate GST-LRRK2 along with LRRK2 interacting partners. Middle panel is membrane after probing with anti-HDAC6 antibody. Upper panel is re-probed with anti-LRRK2 antibody (MJFF2) with HDAC6 signal still visible (*). Lower panel is Ponceau-S signal to show presence of GST. C) Myc-LRRK2-RCKW, RCK, Roc-COR, Roc, COR and kinase domain constructs are indicated. Myc-LRRK2 was immunoprecipitated from HEK293 cells co-transfected with myc-LRRK2 domain constructs and either empty vector (–) or HDAC6. Input and immune pellets were probed with anti-myc and anti-HDAC6 antibodies. D) FLAG-HDAC6 full-length, HD1+2, HD1 or HD2 domain constructs are indicated. FLAG-HDAC6 was immunoprecipitated from HEK293 cells co-transfected with FLAG-HDAC6 full-length, HD1+2, HD1 or HD2 domain constructs with either empty vector or myc-LRRK2-RCK. Input and immune pellets were probed with anti-myc and anti-FLAG antibodies.

To determine if HDAC6 and LRRK2 could directly interact we performed GST pulldown assays in which we incubated a GST-tagged LRRK2 fragment (aa 970-2527) with recombinant His-tagged HDAC6. HDAC6 was readily pulled down with GST-LRRK2 but not with the GST control (Fig. 2B). Because both HDAC6 and LRRK2 have been shown to interact with tubulin, we verified the absence of tubulin in the reaction (SFig. 2).

We next determined the domains of LRRK2 and HDAC6 involved in their interaction in co-immunoprecipitation assays from HEK293 cells (Fig. 2C and D). We found that HDAC6 efficiently interacted with the Roc domain in these assays, and to a much lesser extend with the COR domain; the interaction did not require the ARM/Ankyrin repeat, LRR, or WD40 domains. The kinase domain per se did not interact with HDAC6 (Fig. 2C). Interestingly, while the Roc-COR-Kinase fragment interacted to a similar extend as the Roc domain, interaction of HDAC6 with Roc-COR fragment was markedly reduced and more similar to the COR domain only fragment. Thus, the COR domain appears to impair efficient binding of HDAC6 to the Roc domain, and this can be overcome by the presence of the kinase domain.

HDAC6 interacted with the LRRK2 Roc-COR-Kinase fragment via the histone deacetylase domains and both domains bound LRRK2 to a similar extend. The ubiquitin-binding domain (ZnF-UBP) of HDAC6 was not required to bind the LRRK2 Roc-COR-Kinase fragment, confirming that the interaction was not due to HDAC6 binding to ubiquitinated LRRK2 (Fig. 2D).

We next performed in vitro phosphorylation assays combined with mass spectrometry to identify possible LRRK2 phospho-sites in HDAC6. LRRK2 was found to phosphorylate HDAC6 serine-22 (Fig. 3A). To verify this phosphorylation in cells we co-expressed HDAC6 and wild type LRRK2 or kinase dead LRRK2 D1994A ^17^ in HEK293 cells and determined HDAC6 phospho-serine-22 (pSer-22) levels using phospho-specific antibodies (Fig. 3). Consistent with LRRK2-mediated phosphorylation of expressed HDAC6, pSer-22 increased upon overexpression of wild type but not kinase dead LRRK2 (Fig. 3B). Similarly, we found that overexpression of wild type LRRK2 increased phosphorylation of endogenous HDAC6 (Fig. 3C). Thus, HDAC6 interacts with and is a substrate of LRRK2.

**Figure 3.**
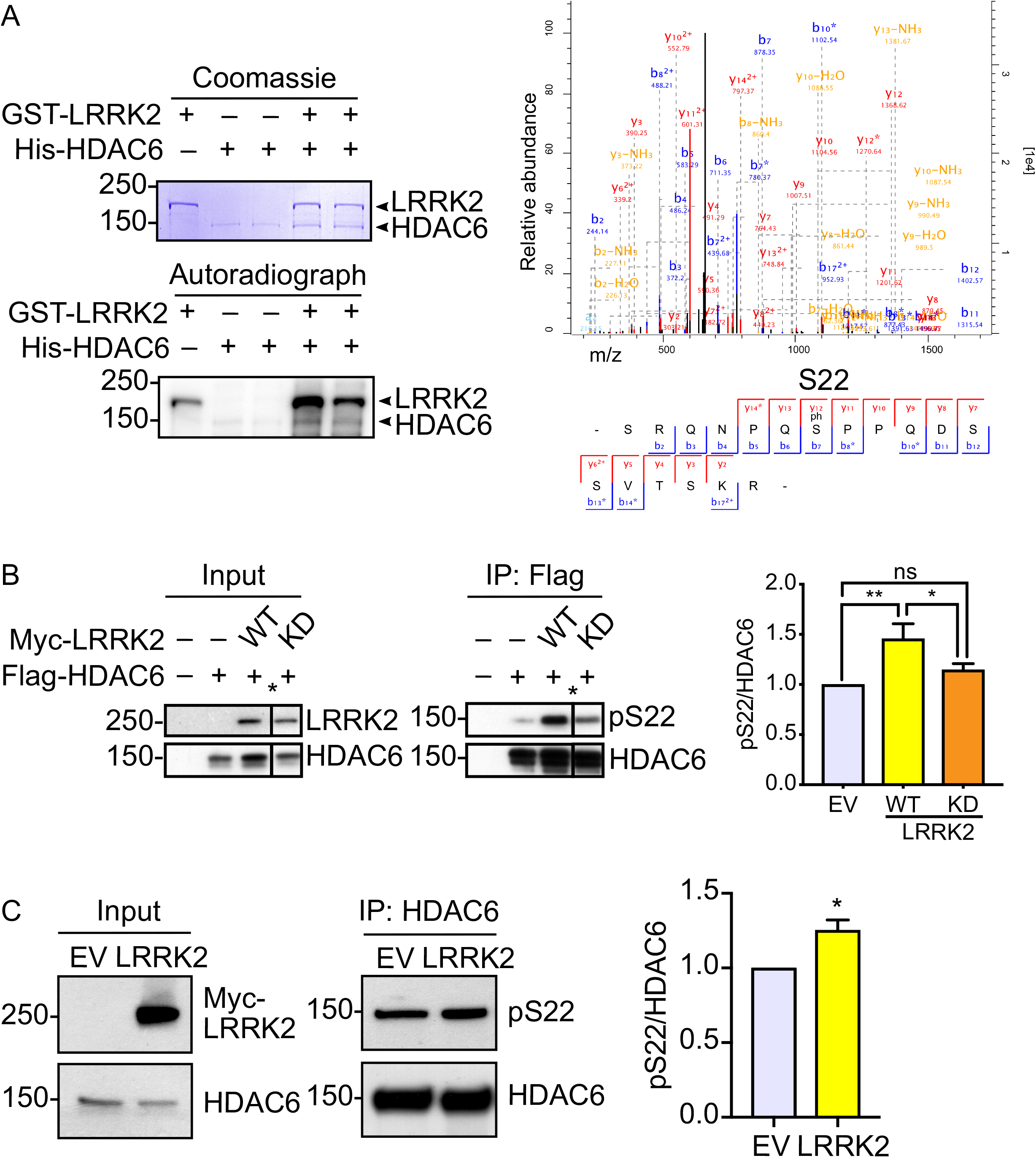
LRRK2 phosphorylates HDAC6. A) Top panel: GST-LRRK2 (aa 970-2527; 500 ng) was incubated with His-HDAC6 (500 ng) and in presence of ^32^P-ATP (57 nM) for 1 h at 37°C. Following separation on SDS-PAGE, proteins were visualised using Coomassie staining and incorporation of ^32^P onto HDAC6 and LRRK2 was detected using a phosphoimager. Bottom panel: Tandem MS/MS spectra of the HDAC6 phosphopeptide RQNPQSPPQDSSVTSK fragment ion series, after collision-induced dissociation. GST-LRRK2 (aa 970-2527; 400 ng) was incubated with His-HDAC6 (1 μg) and 1 mM ATP. Following separation on SDS-PAGE, proteins were digested using trypsin and peptide fragments analysed using LC-MS/MS. High-confidence HDAC6 phosphorylation sites were identified at serine-22 after incubation with LRRK2. B) FLAG-HDAC6 was immunoprecipitated from HEK293 cells co-transfected with FLAG-HDAC6 and empty vector (–), myc-LRRK2-WT or D1994A (KD). Input and immune pellets were probed with anti-myc, HDAC6 and HDAC6 phospho-serine-22 antibodies. An unrelated sample was removed from the scanned blot (*). Quantification shows the ratio of pS22 to total FLAG-HDAC6 in the immune pellets normalised to FLAG-HDAC6 only control (mean ± SEM; one-way ANOVA with Fisher’s LSD test, N = 4 experiments). C) Endogenous HDAC6 was immunoprecipitated from HEK293 cells transfected with empty vector (EV) or myc-LRRK2. The input was probed using anti-myc and anti-HDAC6 antibodies and the immune pellets for total HDAC6 and phospho-serine-22 HDAC6. Quantification shows the ratio of pS22 to total HDAC6 in the immune pellets normalised to empty vector control (data shown as mean ± SEM; Student’s t-Test, N = 3 experiments).

### LRRK2-mediated regulation of aggrephagy requires HDAC6 serine-22 phosphorylation

The above data suggested that LRRK2 mediated phosphorylation of HDAC6 may regulate aggresome formation.

We first checked if LRRK2 kinase activity was required for HDAC6 aggresome formation by rescue of LRRK2 siRNA-treated HEK293 cells with wild type LRRK2 or kinase dead LRRK2 D1994A. While as above wild type LRRK2 efficiently rescued LRRK2 siRNA treatment, kinase dead LRRK2 D1994A was unable to do so (Fig. 4A). Similar results were obtained in LRRK2 KO MEFS (Fig. 4B). Thus, LRRK2 kinase activity is required for HDAC6-dependent aggresome formation.

**Figure 4.**
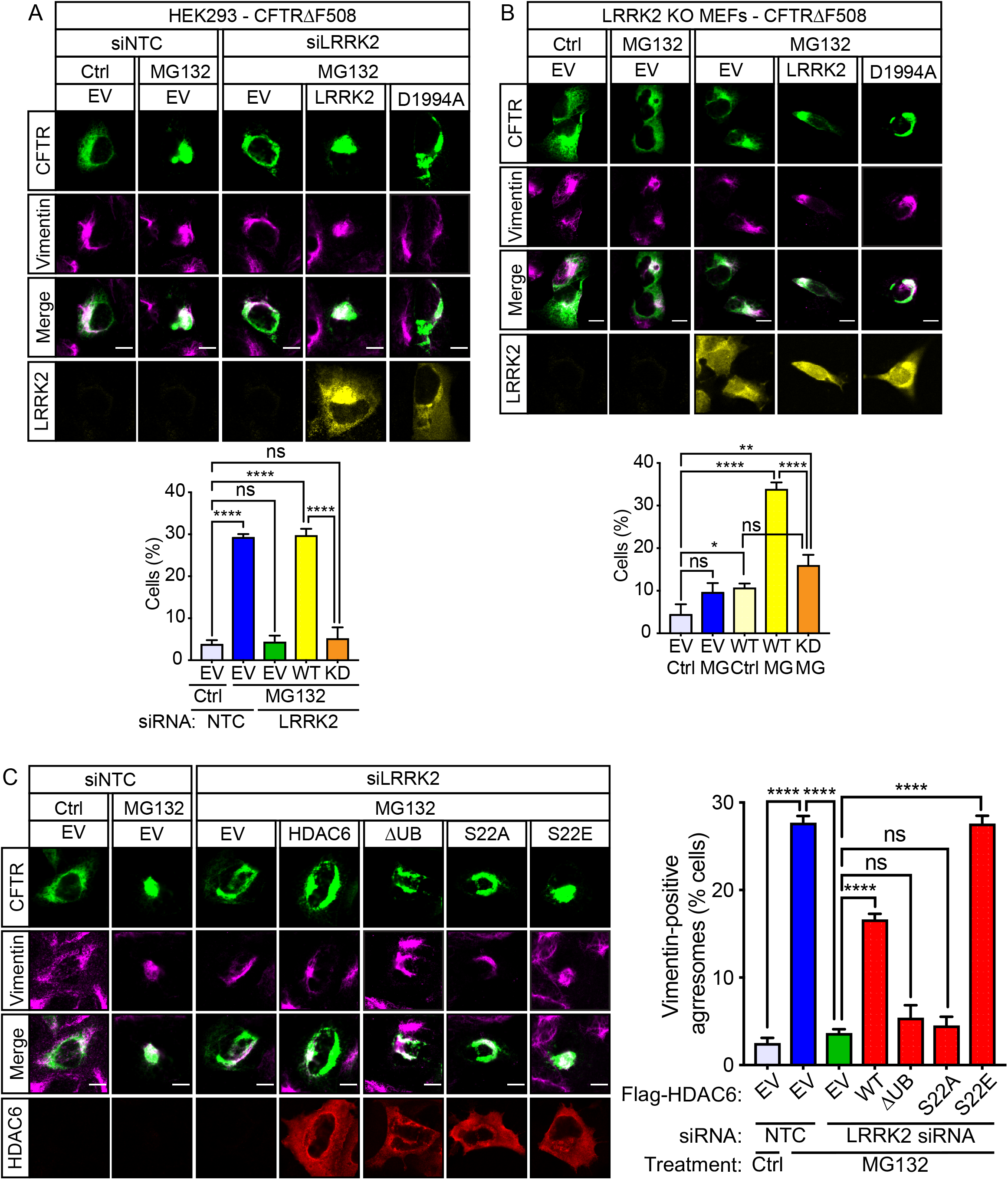
LRRK2-mediated regulation of aggrephagy requires HDAC6 serine-22 phosphorylation. A) Non-targeting (siNTC) or LRRK2 siRNA (siLRRK2)-treated HEK293 cells co-transfected with EGFP-CFTRΔF508 (green) and empty vector (EV), wild type myc-LRRK2 or myc-LRRK2 D1994A (yellow) were treated with vehicle (Ctrl) or MG132 (5 μM) for 4h, fixed and immunostained with an anti-vimentin antibodies (magenta). The percentage of cells containing vimentin-positive EGFP-CFTRΔF508 aggresomes was quantified (mean ± SEM; One-way ANOVA with Fisher’s LSD test, N = 3 experiments, ∼50 cells analysed per condition per experiment). Scale bar, 10 µm. B) LRRK2 KO MEFs co-transfected with EGFP-CFTRΔF508 (green) and empty vector (EV), wild type myc-LRRK2 or myc-LRRK2 D1994A (yellow) were treated with vehicle (Ctrl) or MG132 (5 μM) for 4h, fixed and immunostained with an anti-vimentin antibodies (magenta). The percentage of cells containing vimentin-positive EGFP-CFTRΔF508 aggresomes was quantified (mean ± SEM; One-way ANOVA with Fisher’s LSD test, N = 3 experiments, ∼50 cells analysed per condition per experiment). Scale bar, 10 µm. C) HEK293 cells treated with non-targeting (siNTC) or LRRK2 siRNA (siLRRK2) and co-transfected with EGFP-CFTRΔF508 (green) and empty vector (EV), FLAG-HDAC6-WT, -dUb, -S22A or -S22E (red) were treated with MG132 (5 µM for 4h), fixed and immunostained with anti-FLAG and anti-vimentin (magenta) antibodies. The percentage of cells containing vimentin-positive EGFP-CFTRΔF508 aggresomes was quantified (mean ± SEM; One-way ANOVA with Fisher’s LSD test, N = 3 experiments, ∼60 cells analysed per condition per experiment). Scale bar, 10 µm.

Since HDAC6 was a substrate of LRRK2 (Fig. 3), we reasoned that HDAC6 was likely to be downstream of LRRK2. To test this, we overexpressed HDAC6 on a LRRK2 deficient background and analysed EGFP-CFTRΔF508 aggresome formation. As expected if HDAC6 was downstream of LRRK2, increasing HDAC6 expression partially rescued loss of LRRK2. HDAC6 lacking its ZnF-UBP domain and thus unable to bind ubiquitinated cargo, did not rescue loss of LRRK2 confirming the specificity of the rescue (Fig. 4C). The partial restoration of aggresome formation by HDAC6 expression indicated the possibility that a LRRK2-dependent step was required for full activity. Therefore, using the same experimental paradigm, we next tested the role of HDAC6 pSer-22 by expressing phospho-deficient S22A or phospho-mimicking S22E forms of HDAC6. HDAC6 S22A did not rescue loss of LRRK2, whereas HDAC6 S22E completely restored EGFP-CFTRΔF508 aggresome formation (Fig. 4C). The latter is consistent with a model in which HDAC6 requires LRRK2 phosphorylation on serine-22 for full activity and aggresome formation.

### LRRK2 mediated HDAC6 serine-22 phosphorylation regulates HDAC6 interaction with cytoplasmic dynein

Phosphorylation of HDAC6 on serine-22 has previously been associated with increased deacetylase activity ^18^. Since HDAC6 deacetylase activity is required for aggresome formation ^1^ we compared the deacetylase activity of HDAC6 S22A to that of wild type HDAC6 by quantifying acetylated tubulin on immunoblot and by immunofluorescence. Expression of both HDAC6 S22A and wild type HDAC6 markedly decreased the levels of acetylated tubulin. The effect of HDAC6 S22A was equivalent to wild type HDAC6 in these assays, making it unlikely that its effect on aggresome formation is due to loss of deacetylase activity (Fig. 5A and B).

**Figure 5.**
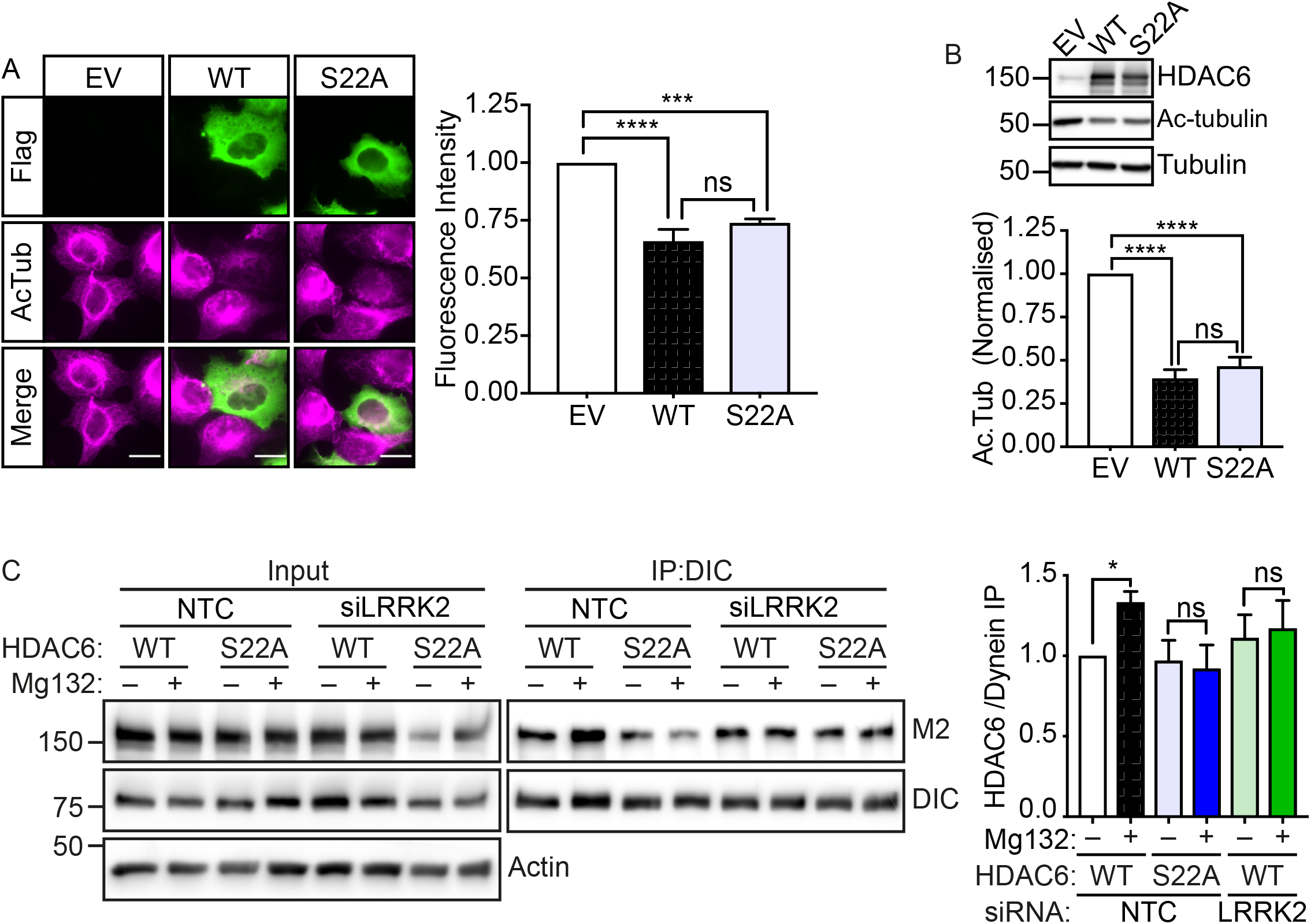
LRRK2 mediated HDAC6 serine-22 phosphorylation regulates HDAC6 interaction with cytoplasmic dynein. A) HEK293 cells were transfected with empty vector (EV), wild type FLAG-HDAC6 or FLAG-HDAC6 S22A, fixed and immunostained with anti-acetylated tubulin and anti-FLAG antibodies. The fluorescence intensity of acetylated tubulin (mean grey value) was quantified per cell and normalised to EV controls (mean ± SEM; One-way ANOVA with Fisher’s LSD test, N = 3 experiments, ∼40 cells analysed per condition per experiment). Scale bar, 10 µm. B) Lysates of HEK293 cells transfected with empty vector (EV), wild type FLAG-HDAC6 or FLAG-HDAC6 S22A were probed using anti-HDAC6, anti-acetylated tubulin, and total tubulin antibodies. Acetylated tubulin was quantified by densitometry and normalised to total tubulin (mean ± SEM; One-way ANOVA with Fisher’s LSD test, N = 7 experiments). C) Cytoplasmic dynein was immunoprecipitated using anti-Dynein IC74 antibody from HEK293 cells treated with non-targeting (siNTC) or LRRK2 siRNA (siLRRK2) and transfected with wild type FLAG-HDAC6 or FLAG-HDAC6 S22A that were treated with vehicle (–) or MG132 (5 µM, 4h). The input and immune pellets were probed using anti-FLAG (M2), and anti-Dynein IC74 antibodies. Actin was used as a loading control. The ratio of FLAG-HDAC6 to dynein intermediate chain (DIC) levels in the immune pellets were determined and normalised to the NTC wild type HDAC6 vehicle control. (mean ± SEM; One-way ANOVA with Fisher’s LSD test, N = 3 experiments).

The binding of HDAC6 to cytoplasmic dynein is crucial for the delivery of ubiquitinated cargo to the aggresome ^1,19^. To test if HDAC6 pSer-22 affects its ability to interact with cytoplasmic dynein, we next inhibited the proteasome in HEK293 cells to induce aggresome formation and evaluated the interaction of wild type HDAC6 and HDAC6 S22A with endogenous cytoplasmic dynein. Induction of aggresome formation increased the interaction of wild type HDAC6 with cytoplasmic dynein as was previously reported (Fig. 5C) ^1^. In contrast, the interaction of HDAC6 S22A with cytoplasmic dynein did not increase after induction of aggresome formation (Fig. 5C). Thus, HDAC6 pSer-22 appears to be instrumental in recruitment of HDAC6 to cytoplasmic dynein after induction of the aggresome pathway. To confirm if the increase in HDAC6 serine-22 phosphorylation and resulting recruitment to cytoplasmic dynein depended on LRRK2 kinase activity we knocked down LRRK2. In absence of LRRK2, wild type HDAC6 was no longer recruited to cytoplasmic dynein after induction of aggresome formation (Fig. 5C).

### Aggresome formation is impaired in LRRK2 PD

LRRK2 G2019S is the most common genetic cause of Parkinson’s disease ^6^. To test if the G2019S mutation affects LRRK2 function in aggresome formation we reconstituted LRRK2 deficient HEK293 cells with LRRK2 G2019S and monitored CFTRΔF508 aggresome formation after proteasome inhibition. While wild type LRRK2 fully restored aggresome formation, LRRK2 G2019S only partially did, indicating that the G2019S mutation reduces LRRK2’s ability to mediate aggresome formation in response to accumulation of ubiquitinated proteins (Fig. 6A). As also noted above in LRRK2 deficient cells (Fig. 1), LRRK2 G2019S did not prevent the formation of vimentin cages per se, but rather appeared to affect the recruitment of CFTRΔF508 to aggresomes.

**Figure 6.**
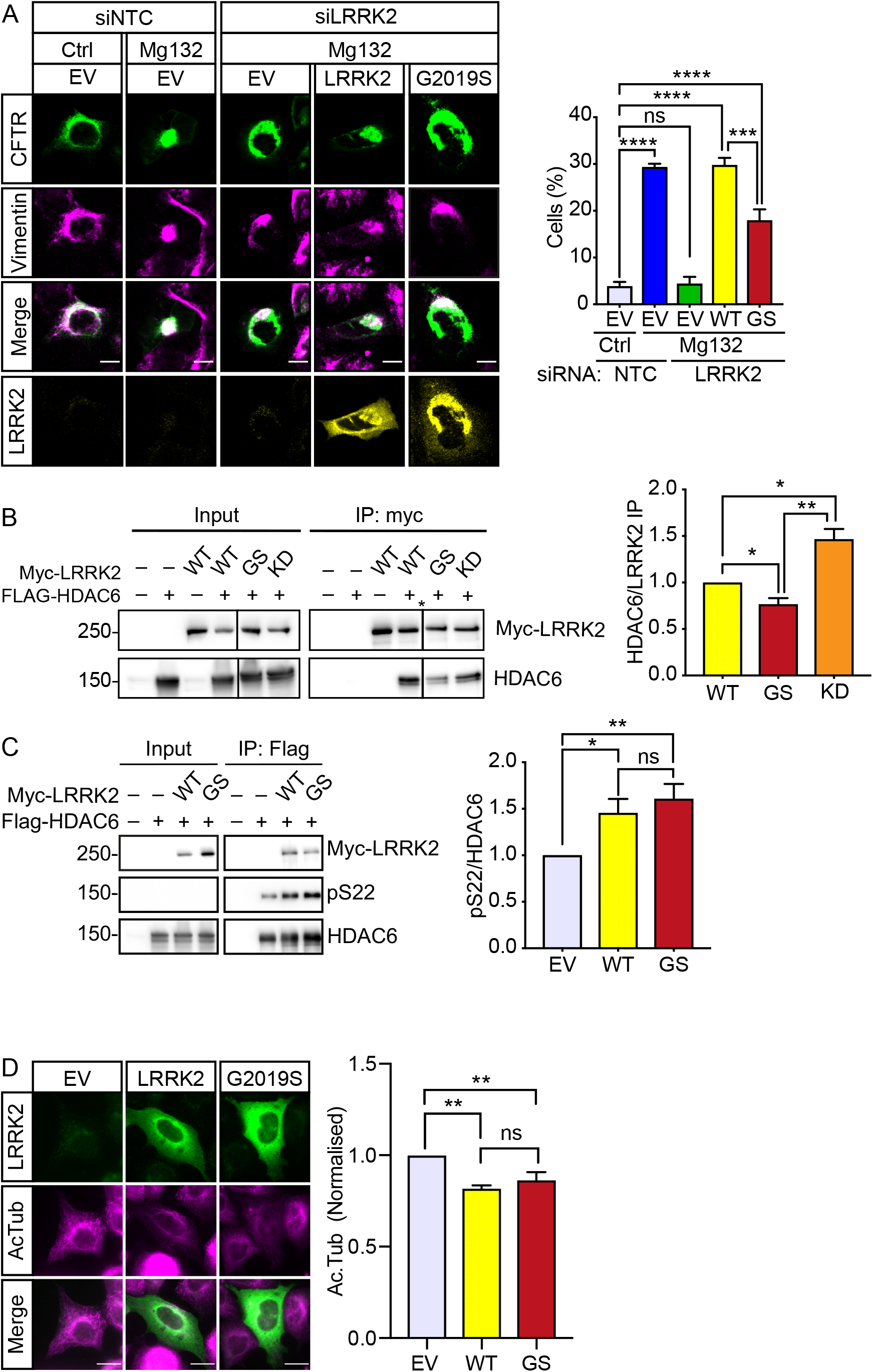
The G2019S mutation in LRRK2 impairs aggresome formation. A) HEK293 cells treated with non-targeting (siNTC) or LRRK2 siRNA co-transfected with EGFP-CFTRΔF508 (green) and myc-LRRK2 wild type or G2019S (yellow). After treatment with vehicle or MG132 (5 µM, 4 h) cells were fixed and immunostained with anti-myc and anti-vimentin (magenta) antibodies. The percentage of cells containing vimentin-positive EGFP-CFTRΔF508 aggresomes was quantified (mean ± SEM; One-way ANOVA with Fisher’s LSD test, N = 3 experiments, ∼4-50 cells analysed per condition per experiment). Scale bar, 10 µm. B) Myc-LRRK2 was immunoprecipitated from HEK293 cells that were co-transfected with empty vector (–) or FLAG-HDAC6 and myc-LRRK2 wild type, G2019S (GS) or D1994A (KD). The immune pellet was probed for LRRK2 and HDAC6 using anti-myc anti-FLAG antibodies. Quantification shows the ratio of HDAC6 to LRRK2 in the immune pellets normalised to wild type LRRK2 (mean ± SEM; one-way ANOVA with Fisher’s LSD test, N = 5 experiments). C) FLAG-HDAC6 was immunoprecipitated from HEK293 cells co-transfected with FLAG-HDAC6 and empty vector (–), myc-LRRK2-WT or G2019S (GS). Input and immune pellets were probed with anti-myc, HDAC6 and HDAC6 phospho-serine-22 antibodies. Quantification shows the ratio of pSer-22 to total FLAG-HDAC6 in the immune pellets normalised to FLAG-HDAC6 only control (mean ± SEM; one-way ANOVA with Fisher’s LSD test, N = 4 experiments). D) HEK293 cells transfected with empty vector (EV), wild type myc-LRRK2 or myc-LRRK2 G2019S, fixed and immunostained with anti-acetylated tubulin and anti-myc antibodies. The fluorescence intensity of acetylated tubulin (mean grey value) was quantified per cell and normalised to EV controls (mean ± SEM; One-way ANOVA with Fisher’s LSD test, N = 4 experiments, ∼40-50 cells analysed per condition per experiment). Scale bar, 10 µm.

To further investigate how the G2019S mutation may affect LRRK2-mediated aggresome formation we tested if the G2019S mutation affected the interaction of LRRK2 with HDAC6, HDAC6 serine-22 phosphorylation, or HDAC6 deacetylase activity. We found that LRRK2 G2019S interacted significantly less with HDAC6 compared to wild type LRRK2 in co-immunoprecipitation assays (Fig. 6B). In contrast, the kinase dead LRRK2 D1994A mutant bound stronger to HDAC6 in these assays (Fig. 6B). Thus, LRRK2 kinase activity inversely correlated with HDAC6 binding. The G2019S mutation did not significantly increase HDAC6 phosphorylation compared to wild type LRRK2, even though numerous groups have shown that the G2019S mutation markedly increases LRRK2 kinase activity (Fig. 6C). Possibly the diminished interaction of LRRK2 G2019S reduces phosphorylation efficiency. Finally, LRRK2 G2019S and wild type LRRK2 stimulated HDAC6 deacetylase activity to the same extend (Fig. 6D). Thus, we propose that decreased interaction of LRRK2 G2019S with HDAC6 reduces the efficiency of serine-22 phosphorylation, and this impairs LRRK2-mediated aggresome formation.

### Aggresome formation is impaired in LRRK2 G2019S patient-derived iAstrocytes

To test if endogenous LRRK2 G2019S impairs aggresome formation in a disease-relevant model we turned to LRRK2 G2019S patient-derived iAstrocytes ^20^. We treated two matched non-disease control and two patient-derived iAstrocyte lines with proteasome inhibitor and monitored endogenous aggresome formation by immunofluorescence microscopy of ubiquitin and vimentin. Both controls consistently formed ubiquitin and vimentin positive aggresomes that also contained HDAC6 (Fig. 7). In contrast, in the two patient-derived iAstrocyte lines, aggresome formation was significantly impaired (Fig. 7). In the patient cells, clusters of ubiquitin that colocalised with HDAC6 were observed but no characteristic accumulation in a perinuclear aggresome occurred (Fig. 7). Thus, in agreement with the data above, while HDAC6 was recruited to ubiquitinated proteins, subsequent recruitment of cytoplasmic dynein appears to be impaired in a model of LRRK2 G2019S PD.

**Figure 7.**
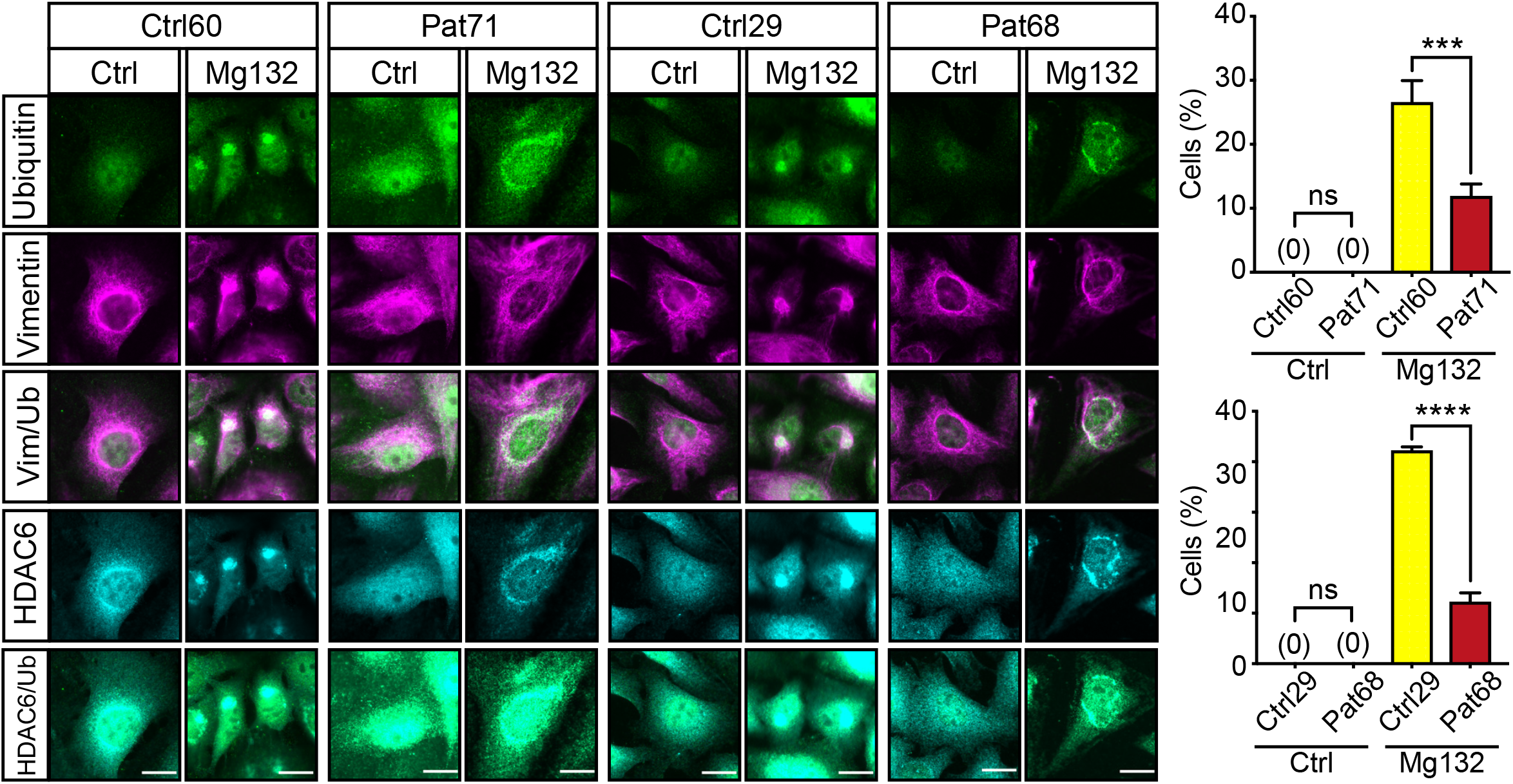
Aggresome formation is impaired in LRRK2 G2019S patient-derived iAstrocytes. LRRK2 G2019S patient-derived iAstrocytes (Pat71, Pat68) and matched controls (Ctrl60, Ctrl29) treated with vehicle or MG132 (5 µM, 14 h) were fixed and immunostained for endogenous ubiquitin (green), vimentin (magenta) and HDAC6 (cyan). The percentage of cells containing vimentin and ubiquitin-positive aggresomes was quantified (mean ± SEM; One-way ANOVA with Fisher’s LSD test, N = 3 experiments, 150-250 cells analysed per condition per experiment). Scale bar, 20 µm.

## Discussion

Accumulation of protein aggregates is a hallmark of many neurodegenerative diseases including PD. Protein homeostasis is maintained by the proteostasis network, a complex regulatory network that controls protein biosynthesis, folding, trafficking, and clearance. Failure of the proteostasis network to deal with misfolded and potentially toxic proteins ultimately is deleterious to cells; this appears to be especially the case for neurons, as exemplified by the common occurrence of aberrant protein folding and aggregate deposition in neurons in neurodegenerative disease ^21^.

Aggregated proteins are mostly removed by autophagy, a process that is also called aggrephagy ^22^. In aggrephagy, ubiquitinated aggregated proteins are recruited to HDAC6 which by binding to cytoplasmic dynein enables the transport of the ubiquitinated protein aggregates along microtubules to the aggresome. The aggresome is insoluble and metabolically stable, and is enclosed by intermediate filaments. The contents of aggresomes may subsequently be degraded by aggrephagy. Thus, the aggresome likely represents a means to sequester aggregated proteins from the cytosol so as to prevent toxicity while awaiting clearance ^23^.

In case of PD, the classic histopathological feature is the Lewy body that comprises mainly ubiquitinated α-synuclein filaments but also numerous other proteins. Lewy bodies share many morphological and biochemical similarities with aggresomes and may form via an aggresome-like mechanism ^2–4^. Several groups have linked HDAC6 to Lewy body formation and have shown that HDAC6 is a constituent of Lewy bodies^1,5,24^. A role for dynactin was also described in Lewy body formation ^25^.

Since our previous studies had linked LRRK2 to HDAC6 ^9^, and LRRK2 itself is present in Lewy bodies ^26^ we hypothesised that LRRK2 may be involved in aggresome formation via HDAC6. Confirming our hypothesis, we found that LRRK2 is required for sequestration of ubiquitinated but not of non-ubiquitinated protein aggregates to aggresomes (Fig. 1). The former is dependent on HDAC6 ^1^ while the latter relies on Bag3 to connect the protein aggregates to cytoplasmic dynein ^27^. We found that HDAC6 binds to LRRK2 and that this interaction is mediated by the HDAC6 deacetylase domains and the LRRK2 Roc domain (Fig. 2). LRRK2 kinase activity appears to regulate HDAC6/LRRK2 interaction (Fig. 2 and 6B). Autophosphorylation of the Roc domain is well established and may regulate GTPase activity and in turn kinase activity ^28–30^. Thus, our data suggests autophosphorylation of the Roc domain may regulate its interaction with HDAC6. Phosphorylation may directly affect binding, or, alternatively, HDAC6/LRRK2 interaction may also be regulated by phosphorylation-induced changes in LRRK2 GTPase activity.

Our data show that LRRK2 phosphorylates HDAC6 on serine-22 (Fig. 3) and that this phosphorylation is crucial for HDAC6-dependent aggresome formation (Fig. 4). Phosphorylation of serine-22 has been reported in a number of large-scale proteomic studies, but its role is not yet clear. It has been suggested that GSK3β-dependent HDAC6 serine-22 phosphorylation increases its deacetylase activity in hippocampal neurons ^18^. However, using a HDAC6 S22A mutant that cannot be phosphorylated, we did not observe an appreciable effect of serine-22 phosphorylation on α-tubulin acetylation levels (Fig. 5A). On the other hand, the same mutant completely failed to rescue aggresome formation (Fig. 4C). Thus, we believe that HDAC6 serine-22 phosphorylation is primarily involved in aggresome formation. Possibly the effect of GSK3β on HDAC6 activity involves other HDAC6 phosphorylation sites.

The interaction of HDAC6 and cytoplasmic dynein increases in response to accumulating levels of misfolded proteins (Fig. 5C) ^1^. Our data show that LRRK2-mediated HDAC6 serine-22 phosphorylation drives HDAC6/dynein complex formation (Fig. 5C). How HDAC6 serine-22 mediates this effect is not yet clear, but since the dynein-binding motif of HDAC6 spans amino acids 439 to 503 ^1^ and thus does not contain serine-22, it is not likely that phosphorylation directly affects the binding interface.

Our data show that LRRK2 is required for aggresome formation (Fig. 1) and that the most common PD-associated LRRK2 mutant, G2019S, did not support aggresome formation to the same extend as wild type LRRK2 (Fig. 6A). This finding seemed counterintuitive because the G2019S mutant has increased kinase activity, and our data show that phosphorylation of HDAC6 by LRRK2 drives aggresome formation (Fig. 5). However, compared to wild type LRRK2 expression of LRRK2 G2019S did not additionally increase HDAC6 phosphorylation (Fig. 6C) while its interaction with HDAC6 was significantly reduced (Fig. 6B). Thus, we suggest that at physiological expression levels, the G2019S mutation behaves as a loss-of-function mutant that reduces LRRK2-mediated phosphorylation of HDAC6 and causes impairment of aggresome formation. In agreement with this, we found a marked defect in aggresome formation in LRRK2 G2019S patient-derived iAstrocytes (Fig. 7).

Inhibition of HDAC6 has been shown to increase α-synuclein levels ^24,31^ and a striking accumulation and aggregation of α-synuclein and ubiquitinated proteins was reported at 20 months of age in LRRK2 KO mice, albeit in kidney ^32^, suggesting that α-synuclein aggregates may be cleared via LRRK2/HDAC6 aggrephagy. Thus, our data supports a model in which reduced aggresome formation in LRRK2 G2019S carriers could lead to increased levels of protein aggregates, including α-synuclein, throughout the cytosol. Consistent with such a model, inhibition of LRRK2 was shown to increase pS129 α-synuclein in LRRK2 G2019S neurons ^33^ and LRRK2 G2019S facilitates α-synuclein neuropathology in aged knock-in mice ^34^. Along the same lines, reduction of LRRK2 protein levels in the brain of mice using LRRK2 anti-sense oligos reduced the amount of fibril-induced α-synuclein inclusions ^35^.

Given the emphasis on LRRK2 kinase inhibitors for PD therapy, further work is warranted to explore LRRK2-mediated regulation of HDAC6-dependent aggresome formation and its relation to α-synuclein pathology.

## Methods

### Plasmids

GFP-CFTRΔF508 expression plasmid was a gift from R. Kopito (Stanford, USA) and GFP-250 was a gift from E. Sztul (University of Alabama, USA). pCMV-Tag-3B-2XMyc-LRRK2-WT and G2019S variants were a gift from M. Cookson (NIH, USA; Addgene #25361 and #25362) ^36^. pCMV-Tag-3B-2XMyc-LRRK2-D1994A was generated by mutagenesis of pCMV-Tag-3B-2XMyc-LRRK2-WT using a QuikChange Lightning Site-Directed Mutagenesis Kit (Agilent Technologies) according to the manufacturer’s instructions. Mutagenic primers were: 5’ acattgtggggtttcagggctcggtatataatcatgg-3’ (forward) and 5’ ccatgattatataccgagccctgaaaccccacaatgt-3’ (reverse). pCMV6-XL4-HDAC6 was obtained from Origene (#SC111132). For cloning of the pCI-neo-3xFLAG-HDAC6 construct, HDAC6 cDNA was generated from pCMV6-XL4-HDAC6 by PCR using Phusion High Fidelity DNA polymerase (NEB) with primers containing restriction sites (SalI-HDAC6-forward: (gtcgac)cccatgctggagtcacct, NotI-HDAC6-reverse: (gcggccgc)ttagtgtgggtggggcata). HDAC6 cDNA was cloned by standard subcloning techniques into pCI-neo-3xFLAG using XhoI/NotI restriction sites. pCI-neo-3xFLAG-HDAC6-dUb, S22A and S22E variants were generated by mutagenesis of pCI-neo-3xFLAG-HDAC6 using primers: 5’ cacagtagacctaatagcaagagagacacacccaat-3’ (Q1150* forward) and 5’ attgggtgtgtctctcttgctattaggtctactgtg-3’ (Q1150* reverse), 5’ cctgagggggcgcctgggggttctg-3’ (S22A forward) and 5’ cagaacccccaggcgccccctcagg-3’ (S22A reverse), 5’ gtcctgagggggctcctgggggttctgc-3’ (S22E forward) and 5’ gcagaacccccaggagccccctcaggac-3’ (S22E reverse), respectively. All constructs were verified by sequencing.

### Cell culture and plasmid transfection

HEK293 and HeLa cells (ATCC) and LRRK2 knock out mouse embryonic fibroblasts (gifted from K. Harvey, UCL, UK) were cultured in Dulbecco’s Modified Eagle Medium (Thermo Scientific) with 4.5 g/l glucose supplemented with 10% fetal bovine serum (FBS; Sigma) and 1 mM sodium pyruvate (Sigma) at 37°C with 5% CO2. Cells were transfected with plasmids using polyethylenimine (PEI) (stock 1 mM; 3 μl/μg plasmid) and used in experiments 24 h post transfection.

### iAstrocyte differentiation

Induced neural progenitor cells (iNPCs) were derived from human skin fibroblasts as previously described ^20^. Punch skin biopsies were taken from two LRRK2 G2019Scarriers and one local control (local ethics obtained, Trondheim, Norway reference 2010/648-4); an additional control fibroblast line was obtained from the Coriell Cell Repository, ND29510.

Human iAstrocyte differentiation was performed as previously described ^20^. Briefly, 50,000 iNPCs were plated in astrocyte differentiation medium (Dulbecco’s Modified Eagle Medium (Thermo Scientific) with 4.5 g/l glucose supplemented with 10% fetal bovine serum (FBS; Sigma)) in a 6-well plate coated with fibronectin (Millipore). Cells were differentiated for 7 days.

### siRNA

Non-targeting control siRNA (MISSION® siRNA Universal Negative Control #1) and LRRK2 siRNA #1 (targeting LRRK2 sequence 5’-ctcgtcgacttatacgtgtaa-3’; ^37^) were purchased from Sigma. LRRK2 siRNA #2 was purchased from ThermoFisher (ID: 263837; ^38^). All LRRK2 knockdowns after the initial validation used both LRRK2 siRNA #1 and #2 in combination. Cells were transfected with siRNA using Lipofectamine RNAiMAX (Invitrogen) according to the manufacturer’s instructions and used in experiments 96 hours post siRNA transfection.

### Treatments

HEK293, HeLa, and iAstrocytes where treated with 5 µM MG132 (Sigma) for 4, 18 and 14 h, respectively. Tubastatin A (Sigma) was used at 10 µM.

### Antibodies

Primary antibodies used for immunoprecipitation (IP), immunoblotting (WB) and immunofluorescence (IF) were as follows: rabbit anti-HDAC6 (D2E5, Cell Signalling, WB: 1:1000, IF: 1:200), rabbit anti-HDAC6 antibody, CT (07-732, Millipore, IF: 1:500) rabbit anti-HDAC6 phospho-S22 (ab61058, Abcam, WB: 1:500), mouse anti-FLAG (M2, Sigma, IP: 1:2000, WB: 1:2000, IF: 1:2000), mouse anti-Myc (9B11, Cell Signalling, IP: 1:2000, WB: 1:2000, IF: 1:2000), rabbit anti-LRRK2 (UDD3, Abcam, IP: 1:1000, WB: 1:1000, IF: 1:100), rabbit anti-LRRK2 phospho-S935 (UDD2, Abcam, WB: 1:1000, IF: 1:200), mouse anti-tubulin (DM1A, Sigma, WB: 1:10,000), mouse anti-ubiquitin (P4D1, Cytoskeleton Inc., IF: 1:250), chicken anti-vimentin (AB5733, Merck, IF: 1:4000). Secondary antibodies used for immunoblotting were horseradish peroxidase-coupled goat anti-mouse and goat anti-mouse IgG (Dako, 1:5000). Secondary antibodies used for immunofluorescence were Alexa fluorophore (488, 568)-coupled goat/donkey anti-mouse IgG, Alexa fluorophore (488)-coupled goat anti-rabbit IgG (Invitrogen, 1:500), BV421-coupled goat anti-rabbit IgG (BD Biosciences, 1:500), Cy5-coupled donkey anti-chicken IgG (Jackson Immuno-Research, 1:500).

### Immunofluorescence

Immunostaining was performed as described previously ^39^. Briefly, HEK293, HeLa cells, LRRK2 knockout MEFs or iAstrocytes grown on glass coverslips were fixed in 3.7% formaldehyde in phosphate-buffered saline (PBS) for 20 min at room temperature. After washing with PBS, residual formaldehyde was quenched by incubation with 0.05 M NH4Cl for 15 min at room temperature. Cells were then permeabilised with 0.2% Triton X-100 in PBS for 3 minutes and washed once with PBS. After fixing, the cells were incubated in PBS with 0.2% fish gelatin (PBS/F) for 30 minutes at room temperature and then with the primary antibody in PBS/F for 1 h at room temperature. After washing with PBS/F, cells were incubated with secondary antibody in PBS/F for 45 min at room temperature and stained with Hoechst 33342 (Thermo Scientific). After a final wash in PBS/F, the samples were mounted in fluorescence mounting medium (Dako).

### Microscopy

Images were recorded using appropriate filtersets (Omega Optical and Chroma Technology) using MicroManager 1.4 software ^40^ on a Zeiss Axioplan2 microscope fitted with a Retiga R3 CCD camera (QImaging), PE-300 LED illumination (CoolLED) and a 63x, 1.4NA Plan Apochromate objective (Zeiss), or a Zeiss Axiovert200 microscope equipped with a Prime sCMOS (Photometrics), PE-4000 LED illumination (CoolLED), and 63x, 1.4NA Plan Apochromate and 100x, 1.3NA Plan Apochromate objectives (Zeiss) and using MetaMorph software (Universal Imaging) on an Olympus IX83 equipped with a Zyla4.2 sCMOS camera (Andor), SpectraX light engine (Lumencor) and OptoLED (Cairn Research) illumination, and 60x, 1.35NA Universal Plan Super Apochromat and 40x, 1.35NA Universal Apochromat objectives (Olympus). Confocal imaging was using a 63x HCX PL APO 1.4 NA oil objective on a Leica TCS SP5 confocal microscope using LAS AF software (Leica Microsystems). All illumination, camera and acquisition settings remained constant during experiments to ensure comparability.

### Image analysis

All image analysis was performed using ImageJ ^41^ and Fiji ^42^. Aggresome counts were performed by quantifying the number of EGFP-CFTRΔF508 or GFP-250-expressing cells exhibiting co-localisation of the GFP signal with vimentin at a single, perinuclear aggresome structure. Where possible, the cells for analysis were selected based on fluorescence in the other channel indicating co-transfection. Acetylated tubulin levels were quantified by measuring the mean grey value in the acetylated tubulin channel. Where possible operators where blinded to the identity of the samples analysed.

### SDS-PAGE and immunoblotting

Cells were washed once with phosphate-buffered saline (PBS), scraped into ice-cold BRB80 buffer (80 mM K-PIPES pH 6.8, 1 mM EDTA, 1 mM MgCl2, 1% (w/v) NP40, 150 mM NaCl, 10 mM NaF, 1 mM Na2VO4, 10 mM β-Glycerophosphate, 5 mM Na_4_P_2_O_7_, and protease inhibitor cocktail (Thermo Scientific)) and lysed on ice for 30 min. Lysates were clarified at 17,000 × g for 30 min at 4°C. Protein concentration was measured by Bradford protein assay following the manufacturer’s protocol (Bio-Rad). SDS-PAGE and immunoblotting were performed as described previously (Webster et. al., 2016). Briefly, samples were separated by SDS-PAGE and transferred to a 0.45 μm nitrocellulose (GE Healthcare) or 0.45 μm PVDF (Merck) membrane by electroblotting (Bio-Rad). Membranes were blocked for 1 h at room temperature in 5% non-fat dry milk (Marvel) or 5% BSA (Sigma) in Tris-buffered saline (TBS) with 0.2% Tween 20. Membranes were incubated with primary antibody in blocking buffer for 1 h at room temperature or overnight at 4°C. Membranes were washed 3 times for 10 min in TBS with 0.2% Tween 20 before incubation with secondary antibodies in blocking buffer for 1 h at room temperature. Membranes were washed a further 3 times for 10 min in TBS with 0.2% Tween 20 and prepared for chemiluminescent signal detection using SuperSignal West Pico chemiluminescent substrate (Thermo Scientific) according to the manufacturer’s instructions. Signals were detected using a GBox chemiluminescence imager (Syngene) or Amersham ECL hyperfilm (GE Healthcare) and quantified using ImageJ (Abramoff et. al., 2004).

### Immunoprecipitation

Cells were washed once with phosphate-buffered saline (PBS), scraped into ice-cold BRB80 buffer (80 mM K-PIPES pH 6.8, 1 mM EDTA, 1 mM MgCl2, 1% (w/v) NP40, 150 mM NaCl, 10 mM NaF, 1 mM Na2VO4, 10 mM β-Glycerophosphate, 5 mM Na_4_P_2_O_7_, and protease inhibitor cocktail (Thermo Scientific)) and lysed on ice for 30 min. Lysates were clarified at 17,000 × g for 30 min at 4°C. 2 mg of protein was incubated with 2-4 μg of primary antibody for 16 h at 4°C. The antibody was captured using 30 μl of 50% Protein G Sepharose beads (Sigma) for 2 h at 4°C. After centrifuging at 3000 × g for 30 s, immune pellets were washed 5 times in ice-cold BRB80 buffer. Protein was eluted in 2x Laemmli buffer and samples were run on SDS-PAGE and immunoblot.

### In vitro binding assay

GST-tagged human LRRK2 wild type, R1441C and G2019S protein (amino acids 970-2527; Invitrogen) was incubated with recombinant His-tagged human HDAC6 protein (EMD Millipore) in 250 μl RB100 buffer (25 mM HEPES pH 7.5, 100 mM KOAc, 10 mM MgCl2, 1 mM DTT, 0.05% (w/v) Triton X-100, 10% (v/v) glycerol) for 1 h at 4°C. GST-tagged proteins were captured by incubation with 20 μl glutathione sepharose (GE Healthcare) beads for 30 min at 4°C. Following centrifugation for 1 min at 2000 × g, beads were washed 3 times with RB100 buffer and protein was eluted in 20 μl glutathione elution buffer (50 mM Tris-HCl pH 7.5, 100 mM NaCl, 40 mM reduced glutathione) for 10 min at room temperature. Samples were run on SDS-PAGE and immunoblot.

### In vitro kinase assay

GST-tagged human LRRK2 wild type, R1441C and G2019S protein (amino acids 970-2527; Invitrogen) and His-tagged human HDAC6 protein (EMD Millipore) were incubated with 57 nm 32P-ATP in kinase buffer (50 mM Tris HCl, 10 mM MgCl2, 1.5 mM 2-mercaptoethanol, 150 mM NaCl) for 1 h at 37°C. Samples were run on SDS-PAGE and proteins were stained with Coomassie Brilliant Blue R-250 (Thermo Scientific). The gel was dried onto Whatman paper in a 583 vacuum dryer (Bio-Rad) at 80°C for 1 h. 32P radiolabel was detected using a PMI autoradiograph imager (Bio-Rad).

### Mass spectrometry

GST-LRRK2 (aa 970-2527, Invitrogen) was incubated with His-HDAC6 (EMD Millipore) and 1 mM ATP in kinase buffer for 1 h at 37°C. 20 mM EDTA was added to halt the reaction. Samples were boiled in NuPAGE LDS sample buffer (Thermo Scientific) for 10 min at 70°C in a Thermomixer (Eppendorf) at 800 rpm. Samples were incubated with 10 mM iodoacetamide for 30 min at room temperature in the dark and separated on SDS-PAGE. Proteins were stained with Brilliant Blue G Colloidal Coomassie (Sigma) and the HDAC6 band of interest was excised. Excised gel slices were incubated with 50% acetonitrile/50mM ammonium bicarbonate for 2 h at room temperature or 4°C overnight with gentle shaking at 600 rpm in a Thermomixer (Eppendorf), before being incubated in 100% acetonitrile for 15 min at room temperature. For trypsin digestion, gel slices were incubated in 1 ng/μl trypsin (Thermo Scientific) in 50 mM ammonium bicarbonate for 1 h at 37°C with shaking at 600 rpm in a Thermomixer followed by incubation for 16 h at room temperature.

Peptides were extracted in 100% acetonitrile for 15 min at 37°C with shaking at 600 rpm in a Thermomixer, before transfer of the supernatant to a new 1.5 ml peptide collection tube and incubation of the gel slices in 0.5% formic acid for 15 min at 37°C with shaking at 600 rpm in a Thermomixer. The gel slices were further incubated with 100% acetonitrile for 15 min at 37°C with shaking at 600 rpm in a Thermomixer and the supernatant was removed and added to the previous peptide collection tube. This process was repeated once with 0.5% formic acid followed by two times with 100% acetonitrile, and the final peptide collection tube was dehydrated in a SpeedVac (Thermo Scientific) for 16 h at room temperature. The resulting peptides were stored at −20°C.

Samples were analysed using LC-MS/MS on an Ultimate 3000 RSLC Nano LC System (Dionex) coupled to an LTQ Orbitrap Elite hybrid mass spectrometer (Thermo Scientific) equipped with an Easy-Spray (Thermo Scientific) ion source. Peptides were desalted online using an Acclaim PepMap100 capillary trap column (Thermo Scientific) and separated using 120 min RP gradient (4-30% acetonitrile/0.1% formic acid) on an Acclaim PepMap100 RSLC C18 analytical column (Thermo Scientific) with a flow rate of 0.25 μl/min. The mass spectrometer was operated in standard data dependent acquisition mode controlled by Xcalibur software (Thermo Scientific). The instrument was operated with a cycle of one MS (in the Orbitrap) acquired at a resolution of 60,000 at m/z 400, with the top 20 most abundant multiply-charged (2+ and higher) ions in each chromatographic window being subjected to CID fragmentation in the linear ion trap. An FTMS target value of 1e6 and an ion trap MSn target value of 5000 were used. Dynamic exclusion was enabled with a repeat duration of 30 s, an exclusion list of 500 and an exclusion duration of 60 s. Lock mass of 401.922 was enabled for all experiments.

Data was analysed using MaxQuant software ^43^. Data was searched against a UniProt human sequence database using the following search parameters: trypsin with a maximum of 2 missed cleavages, 7 ppm for MS mass tolerance, 0.5 Da for MS/MS mass tolerance, with acetyl (Protein N-term), phospho (STY) and oxidation (M) as variable modifications and carbamidomethyl (C) as a fixed modification. A protein FDR of 0.01 and a peptide FDR of 0.01 were used for identification level cut offs and high confidence phosphorylation sites were defined using a PEP cut-off of 0.01. Class I phosphorylation sites were defined with a localization probability of >0.75 and a score difference of >5.

### Statistical analysis

All calculations were performed using Excel (Microsoft Corporation, Redmond, WA) and statistical analysis was performed in Prism 7 and 8 (GraphPad Software Inc., San Diego, CA). Statistical significance was determined by one-way analysis of variance (ANOVA) followed by Fisher’s least significant difference post hoc test or by t-test as indicated; *= P ≤ 0.05, **= P ≤ 0.01, ***= P ≤ 0.001, ****= P ≤ 0.0001.

## Acknowledgements

We thank Kirsten Harvey, Ron Kopito, Elizabeth Sztul and Mark Cookson for generously sharing reagents, and all the members of the De Vos and Grierson labs at SITraN for helpful discussions. This work was supported by the following grants: UK Medical Research Council (MR/K005146/1 to K.J.D.V.), Parkinson’s UK Senior Fellowship (F-1301 to H.M.), Parkinson’s UK small grant (K-1506 to L.F. and H.M.), Academy of Medical Sciences/Wellcome Trust Springboard Award (SBF002\1142 to L.F.), and Rosetree’s Trust M580 (H.M. and R.M.). Funding to pay the Open Access publication charges for this article was provided by the UK Medical Research Council. This research was in part supported by the NIHR Sheffield Biomedical Research Centre

## Author contributions

R.M.L, C.S.B, K.C. performed experiments, analysed data and made figures.

A.S., R.M., L.F., and H.M. generated iAstrocyte cultures.

M.O.C. performed mass spectrometry experiments and analysed data.

J.O.A. and G.B. provided LRRK2 G2019S fibroblasts and controls.

K.J.D.V. conceived the study, performed experiments, analysed data, made figures and wrote the manuscript with input of all authors.

## Materials & Correspondence

Correspondence and material requests should be addressed to K.J.D.V

## Data availability

All data files and files produced for statistical analysis are available on request.

---

**Supplementary Figure 1.**
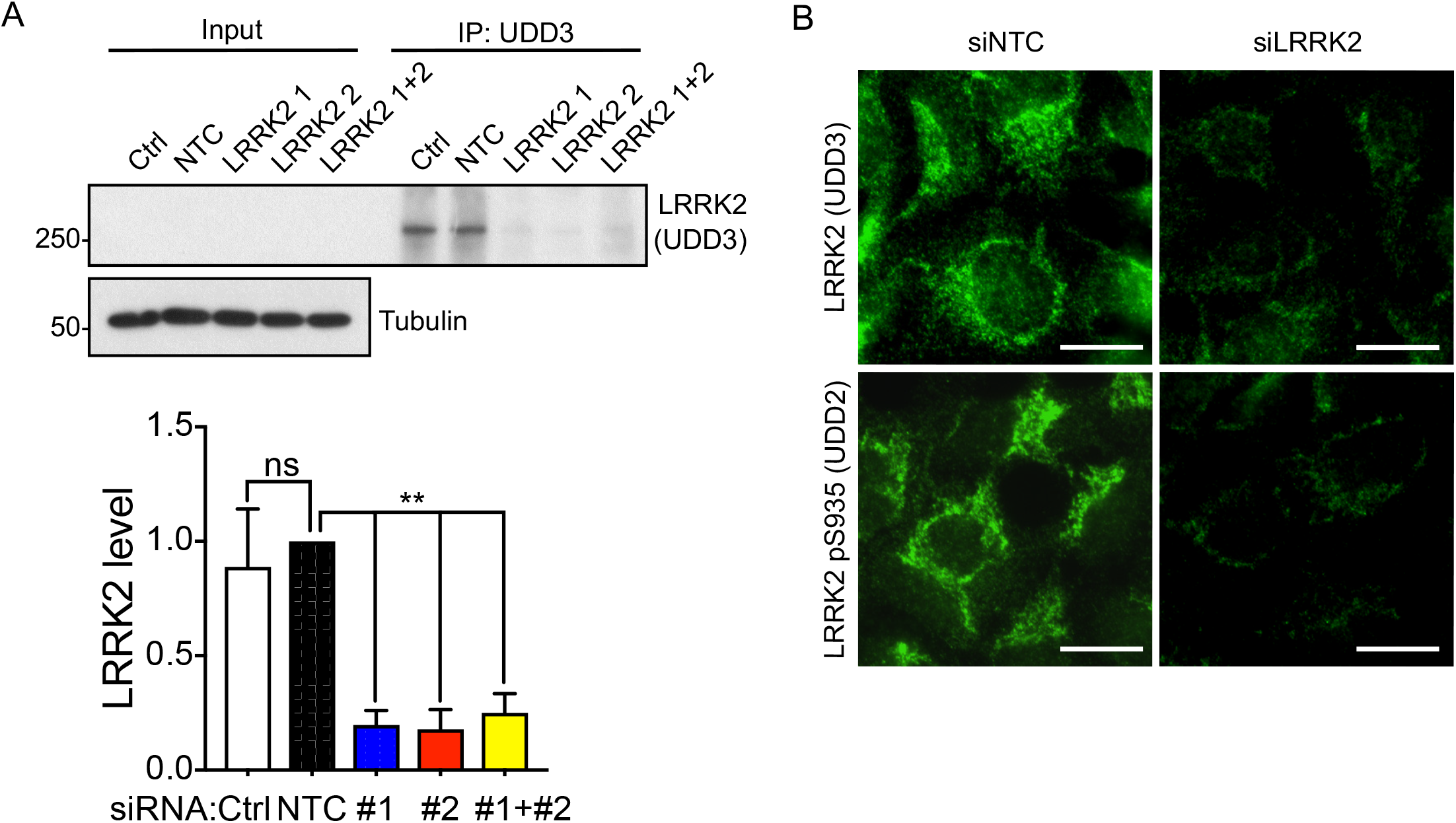
Validation of LRRK2 siRNA knockdown in HEK293 cells. A) LRRK2 was immunoprecipitated using anti-LRRK2 (UDD3) antibodies from HEK293 cells treated with vehicle (Ctrl), non-targeting control siRNA (siNTC), or LRRK2 siRNA #1, #2 or #1 + #2. Immunoprecipitated LRRK2 was detected on immunoblot using anti-LRRK2 (UDD3). Tubulin was used as loading control. The level of immunoprecipitated LRRK2 was determined by densitometry and normalised to the NTC siRNA sample (mean ± SEM; One-way ANOVA with Fisher’s LSD test, N = 3 experiments). B) HEK293 cells treated with non-targeting control siRNA (siNTC) or LRRK2 #1 + #2 (siLRRK2) were fixed and immunostained using UDD3 and UDD2 anti-LRRK2 antibodies. Scale bar, 20 µm.

**Supplementary Figure 2.**
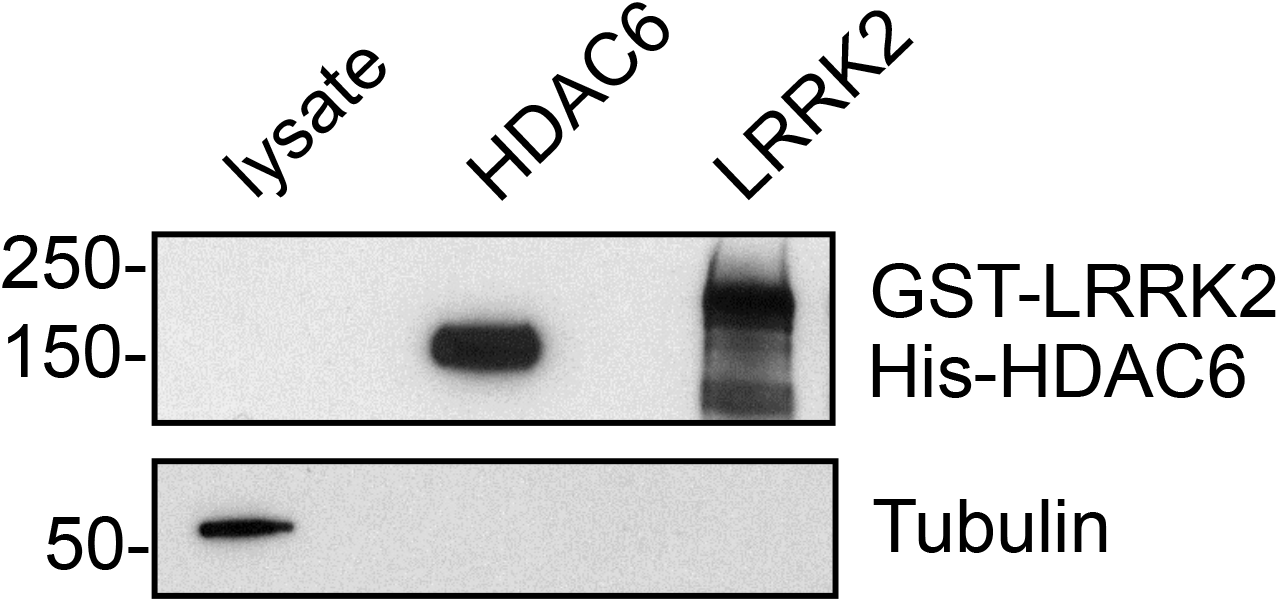
Tubulin is absent from recombinant HDAC6 and LRRK2 preparations. 1 μg His-HDAC6 and GST-LRRK2 variants were tested for presence of tubulin using an anti-tubulin DM1A antibody. HDAC6 and LRRK2 were detected using anti-HDAC6 (D2E5, Cell Signalling) and anti-LRRK2 (MJFF2, Abcam) antibodies respectively. 10 μg total HEK293 cell lysate was used as positive control.

**Supplementary Figure 3.**
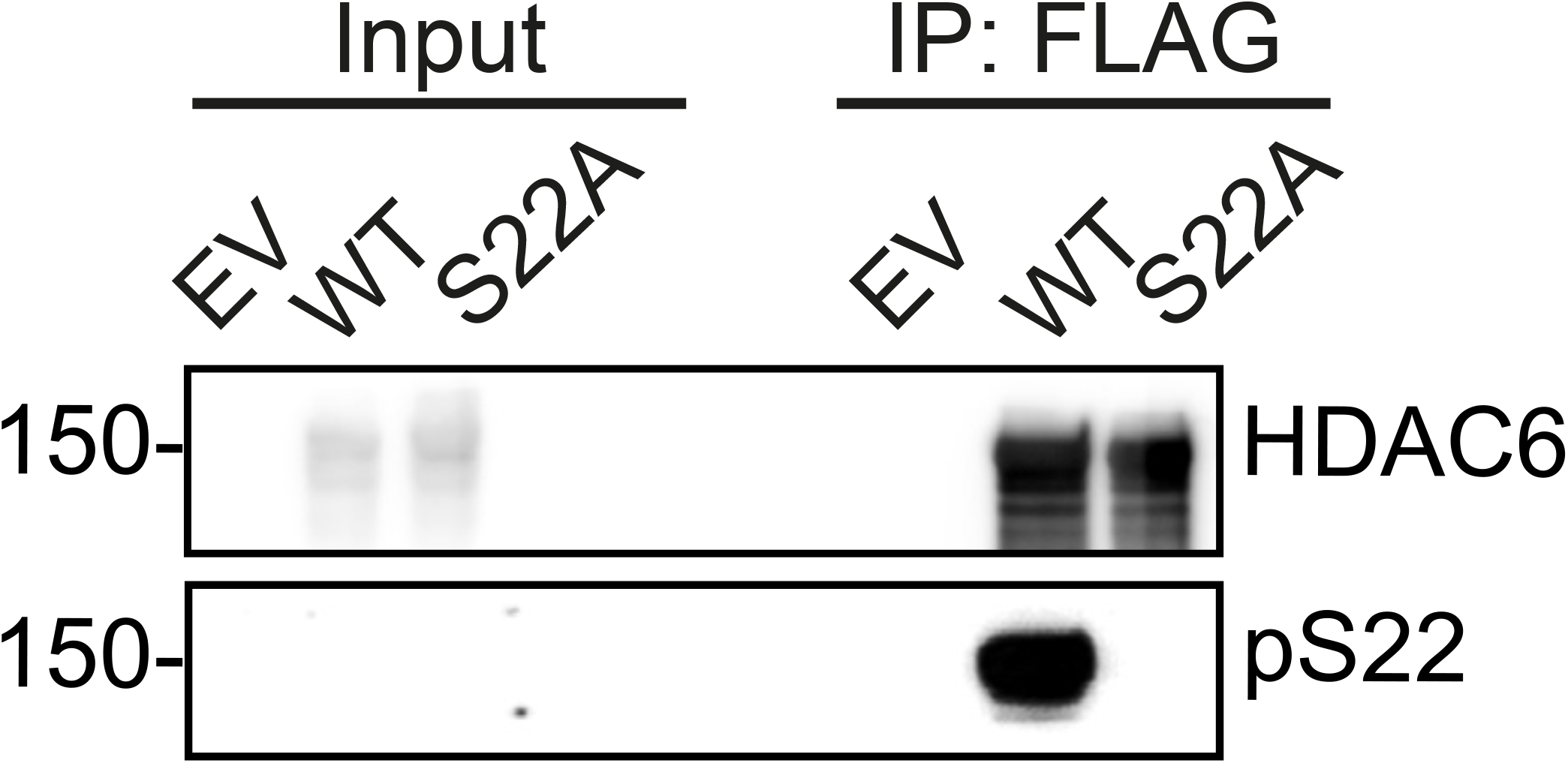
HDAC6 phospho-serine 22 antibody is site-specific. HEK293 cells were transfected with empty vector (EV), FLAG-tagged wild type HDAC6 (WT) or FLAG-HDAC6 S22A (S22A) and HDAC6 was immunoprecipitated using anti-FLAG antibodies. Immune pellets were probed for total HDAC6 and HDAC6 pSer-22.

